# Spatial variability of agricultural soil carbon dioxide and nitrous oxide fluxes: characterization and recommendations from spatially high-resolution, multi-year dataset

**DOI:** 10.1101/2024.08.02.606447

**Authors:** Nakian Kim, Chunhwa Jang, Wendy H. Yang, Kaiyu Guan, Evan H. DeLucia, DoKyoung Lee

## Abstract

Mitigating agricultural soil greenhouse gas (GHG) emissions can contribute to meeting the global climate goals. High spatial and temporal resolution, large-scale, and multi-year data are necessary to characterize and predict spatial patterns of soil GHG fluxes to establish well-informed mitigation strategies, but not many of such datasets are currently available. To address this gap in data we collected two years of high spatial resolution (7.4 sampling points ha^−1^ over 2.0 to 5.4 ha area) in-season soil carbon dioxide (CO_2_) and nitrous oxide (N_2_O) fluxes from three commercial sites in central Illinois, one conventionally managed continuous corn and two under conservation practices in corn-soybean rotations typical of the region. At the field-scale, the spatial variability of CO_2_ was comparable across sites, years, and management practices, but N_2_O was on average 77% more spatially variable in the conventionally managed site. Analysis of N_2_O hotspots revealed that although they represent a similar proportion of the sampling areas across sites (conventional: 12%; conservation: 13%), hotspot contribution to field-wide emission was greater in the conventional site than in the conservation sites (conventional: 51%; conservation: 34%). Also, the spatial patterns, especially hotspot locations, of both gases were inter-annually inconsistent, with hotspots rarely occurring in the same location. Overall, our result indicated that traditional field-scale monitoring with gas chambers may not be the optimal approach to detect GHG hotspots in row crop systems, due to the unpredictable spatial heterogeneity of management practices. Still, our sensitivity analysis on the dataset demonstrated that sampling at a spatial resolution of 1.6 and 5.6 points ha^−1^ can provide reliable (< 25% error) estimates of field-scale soil CO_2_ and N_2_O fluxes, respectively.

## 1. Introduction

Agricultural soils are an important source of greenhouse gas (GHG), responsible for 17% of global agricultural GHG emissions (Lamb et al., 2021). In non-flooded agricultural systems, these emissions mainly comprise of carbon dioxide (CO_2_) from mineralization of soil organic matter, microbial heterotrophic respiration, and crop root autotrophic respiration, and nitrous oxide (N_2_O; 298 CO_2_eq) from nitrification and denitrification of nitrogen (N) fertilizer inputs. Especially, N_2_O from agricultural soils contribute approximately 69% of the total anthropogenic N_2_O (Shukla et al., 2022). Thus, mitigating agricultural soil GHG will help increase the much-needed margin in the tight race toward meeting the global climate goal to limit global warming by 1.5`°`C as mandated in the Paris Agreement (UNFCCC, 2015).

High-quality data from reliable soil gas flux measurements are necessary to advise effective GHG mitigation strategies. However, soil GHG fluxes are spatiotemporally variable, as demonstrated by empirical data (Charteris et al., 2021; Hatfield and Parkin, 2012; Yang et al., 2022) and simulation models (McNunn et al., 2020). Residual errors from spatial heterogeneity are significant sources of variability in soil GHG flux measurements, leading to uncertainties that hinder accurate monitoring of soil GHG fluxes (Charteris et al., 2021, 2020; Kravchenko et al., 2017; Kravchenko and Robertson, 2015; McDaniel et al., 2017). Especially, N_2_O is known for high spatial variability, where hotspots contribute disproportionately to field-scale emissions (Cowan et al., 2015; Turner et al., 2016). Thus, failure to accurately represent these N_2_O flux hotspots can greatly over or underestimate field-scale emissions. Moreover, GHG, especially N_2_O, is temporally dynamic and low temporal resolution datasets lead to undesirable level of errors (Dencső et al., 2021; Taki et al., 2019). Thus, large-scale and high spatiotemporal resolution measurement campaigns that can capture these spatial and temporal variability is necessary to study the soil GHG fluxes.

Characterizing the spatial variability of soil GHG fluxes can help refine sampling strategies, improve quantification methods, and develop tools like predictive models that will help collect datasets more efficiently. For this goal, multiple large-scale, high-resolution campaigns have been conducted, despite their tremendous costs and labor. For example, field-scale soil GHG flux studies have identified soil N level (Cowan et al., 2015), soil moisture (Charteris et al., 2021; Mason et al., 2017), and topography (Ashiq et al., 2021; Turner et al., 2016) as important predictors of N_2_O spatial patterns and hotspots. The spatial resolutions of these studies ranged as low as four measurement points ha^−1^ (Turner et al., 2016) to 55 points ha^−1^ (Charteris et al., 2021), although such high spatial resolution measurement was done only at a single time point during a season. These reports suggest that biogeochemical mechanisms behind soil GHG flux indeed dictate its field-scale spatial pattern. However, the inter-annual stability of these observed spatial patterns remains largely unknown, as there have not been enough number of multi-year datasets. If GHG spatial patterns change each year due to biogeochemical and management factors, measurement protocols and models based on a single year of data can be misleading. Unfortunately, testing this requires multi-year campaigns over consistent locations, whose costs make such datasets even rarer. A two-year campaign on two corn-soybean rotation sites used time stability analysis to identify temporally stable monitoring points to better represent field-scale soil N_2_O fluxes (Ashiq et al., 2021). However, the factors behind the difference between inter-annually consistent versus transient hotspots were not within their scope, nor testing whether the spatial patterns changed yearly. Other than this past study, currently there have not been other multi-year, high-resolution, field-scale measurement campaigns. Thus, the inter-annual stability, especially between growing seasons, of soil GHG flux spatial patterns and its determinants have not been well characterized.

The goal of this study was to collect multi-year, multi-site, field-scale, and whole in-season datasets to characterize the field-scale spatial variability of soil GHG fluxes. For this goal, we carefully selected three on-farm research sites representing typical US Midwestern corn and soybean cropping systems: conventional tillage and continuous corn, compared to the other two sites under reduced or no-till and corn-soybean rotation. Thus, our objectives were to i) characterize the field-scale spatial variability of soil CO_2_ and N_2_O fluxes and compare it across sites and years, ii) evaluate the inter-annual stability of each site’s spatial patterns, especially hot/coldspots, and iii) capitalize on our high-resolution, large-scale dataset to perform sensitivity analysis by simulating lower-resolution datasets to determine the optimal spatial resolution for field-scale soil GHG flux measurement.

## 2. Materials and Methods

### 2.1. Experimental site description

The three experimental sites were in central Illinois (Supplementary Fig. 1), and the size of each site was approximately 35 hectares, which were managed by local farmers who owned each site. Bondville site was located northwest of Bondville, IL (720 ft elevation). This site consisted of 30% Drummer series soil, 30% Flanagan silt loam (mesic Aquic Argiudolls), and 20% Catlin silt loam (mesic Oxyaquic Argiudolls) (USDA-NRCS, 2022). The monitored part of Bondville site was an oval shaped area approximately 216 m wide and 242 m long in 2021, and a rectangular area that was 108 m wide and 106 m long in 2022. Pesotum site was located east of Pesotum, IL (670 ft elevation). Pesotum site is composed of 50% Drummer and 50% Flanagan. For both years, rectangular area of approximately 110 m wide and 148 m long was monitored in this site. Villa Grove site was located north of Villa Grove, IL (650 ft elevation). Villa Grove site was 70% Drummer silt loam clay (mesic Aquic Argiudoll) and 30% Millbrook silt loam (mesic Udollic Endoaqualfs) with a slope of 0 to 2% (Soil survey 2022). The monitored area of this site was approximately 112 m wide and 147 m long in 2021, and 188 m wide and 221 m long in 2022.

### 2.2. Field management practices

Each of the three sites was managed differently. Bondville and Pesotum sites were in corn (2021) and soybean (2022) rotations under conservation tillage practices. In 2021, Bondville site was under no-till while Pesotum site was strip tilled in the fall of 2020. In 2022, Bondville site was under light vertical tillage before planting, and Pesotum site was under no-till. Meanwhile, Villa Grove site was in continuous corn under conventional deep chisel tillage in the fall for both years. The annual total N fertilization rates for corn years were similar across sites at 220 kg N ha^−1^. The application timing (fall application, spring application at planting, and side dressing) and the amounts for each application differed by site. The detailed descriptions of the management practices and their dates are summarized in Supplementary Table 1.

### 2.3. Soil GHG flux measurements

The soil GHG fluxes were measured during two growing seasons, 2021 and 2022, over a large area at each site to determine the inter-annual consistency of their spatial patterns and monitor any effects from different management practices. For all sites, the sampling points were set up in a grid, about 37 m apart horizontally and vertically, so that there were 7.4 sampling points per hectare, each point representing a 0.1 ha area, covering an average of 2.4 ha for each site. However, to collect a more extensive dataset over a larger area within the practical limits of available labor and equipment, one of the sites was selected each year (2021: Bondville; 2022: Villa Grove) to collect data over an expanded sampling area. Hence, in Bondville site, 40 points covered a 5.4 ha area in 2021, and 16 points in a 4 x 4 grid over a 2.0 ha area in 2022. For Pesotum site, in both years, 19 points in 5 x 4 grid covered a 2.7 ha area. In Villa Grove site, 2021, there were 18 points installed over about a 2.8 ha space, which extended to 40 points in 2022 that covered 5.4 ha. The GPS locations of these sampling points were recorded (R8s, Trimble, Westminster, CO, USA) to keep them consistent between years. The arrangements of the sampling points are illustrated in Fig. 1.

**Figure 1.**
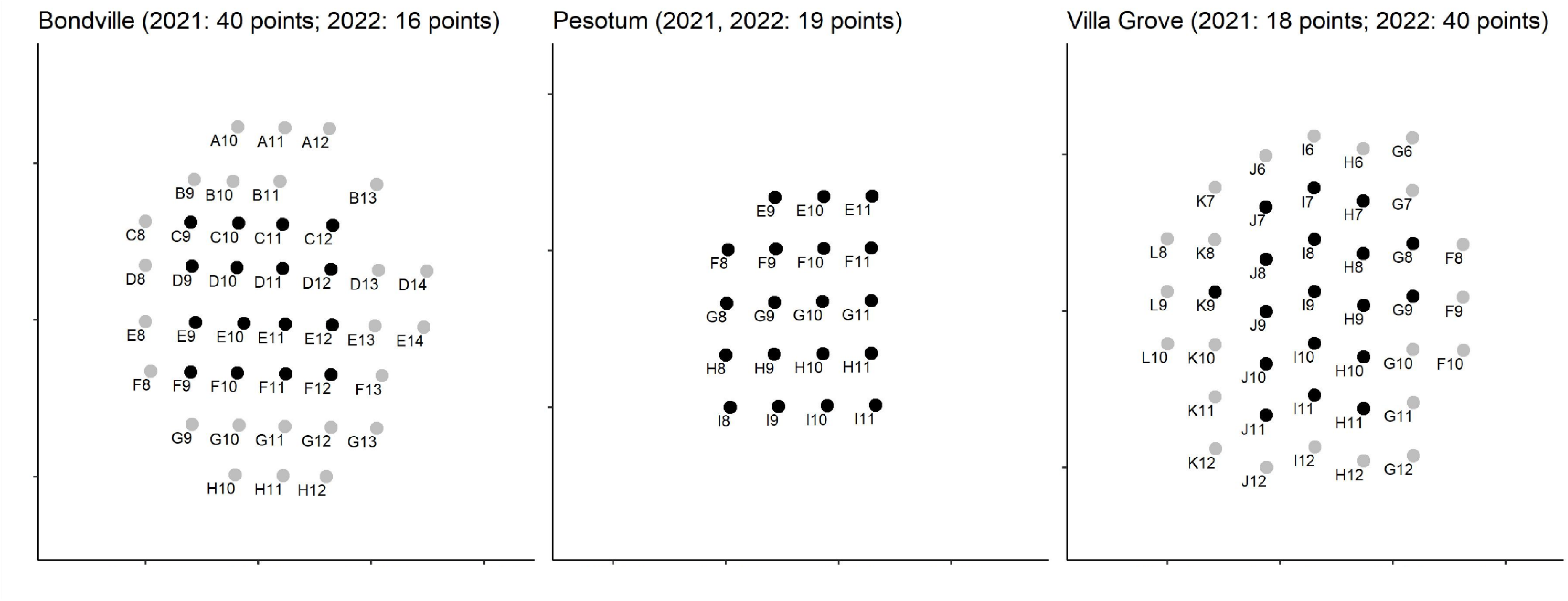
GPS locations of soil GHG flux sampling points of each site. Black points indicate those sampled in both years (2021 and 2022) and grey points indicate those sampled in one of the years (2021 for Bondville and 2022 for Villa Grove) from expanded measurement campaign. The latitude and longitude of these points are not reported for the privacy of the commercial sites.

Soil GHG fluxes were measured manually using closed dynamic surface flux chambers. Polyvinyl chloride (PVC) collars with internal diameter of 20 cm were installed at these points, leaving 5 cm aboveground. Each year after planting, soil CO_2_ and N_2_O fluxes were measured once a week, then once in every two weeks after the last N_2_O fluxes peak following N fertilization, around late June. The durations of in-season measurements for each year and site are shown in Table 1. At each point for each sampling event, a chamber of 4 L in volume connected to portable gas analyzers (DX4040 and GT5000, Gasmet Technologies Oy, Vantaa, Finland) was fitted tightly over the collar, during which the GHG concentrations were measured every thirty (DX4040) or twenty (GT5000) seconds for four minutes. The first minute of a measurement was discarded to account for the leftover gas from the previous measurement. For each measurement, the linear slope of GHG concentration over time and the R^2^ value were calculated for CO_2_ and N_2_O. For CO_2_, if R^2^ rounded to the hundredth was below 0.99, the measurements for both CO_2_ and N_2_O were discarded. For N_2_O, datapoints with Cook’s distance larger than 4.0 divided by the number of datapoints for the respective measurement were removed as outliers unless they were the first or last datapoints; if slope was not statistically significantly different from zero (p-value > 0.100), the measurement was considered a noise and zero flux was recorded (Xia and Wander, 2022). The slope was used to calculate the GHG fluxes in μg cm^−2^ s^−1^, which was then converted to g m^−2^ day^−1^ for CO_2_ and mg m^−2^ day ^−1^ for N_2_O.

**Table 1.** Growing season cumulative soil CO_2_ (g m^−2^) and N_2_O (mg m^−2^) fluxes averaged for nodes of manual chamber measurements. The site, year, the cropping system, the duration of measurements, the number of nodes (n), standard deviation (SD), and coefficient of variations (CV) are also shown.

| Site | Year | Crop | Tillage | Duration | n | mean cumulative<br>CO<sub>2</sub> | SD | CV (%) | mean cumulative<br>N<sub>2</sub>O | SD | CV (%) |
| --- | --- | --- | --- | --- | --- | --- | --- | --- | --- | --- | --- |
| Bondville | 2021 | Corn | No-till | 5/5 - 9/7 | 40 | 1225 | 219 | 18 | 155 | 109 | 70 |
|  | 2022 | Soybean | Light vertical | 5/24 - 9/19 | 16 | 1224 | 280 | 23 | 86 | 81 | 94 |
| Pesotum | 2021 | Corn | Strip | 5/7 - 9/1 | 19 | 1239 | 298 | 24 | 172 | 142 | 83 |
|  | 2022 | Soybean | No-till | 5/24 - 9/19 | 19 | 1148 | 236 | 21 | 70 | 44 | 63 |
| Villa Grove | 2021 | Corn | Deep chisel | 5/7 - 9/1 | 18 | 1511 | 411 | 27 | 604 | 812 | 135 |
|  | 2022 | Corn | Deep chisel | 5/19 - 10/3 | 40 | 1690 | 375 | 22 | 785 | 1092 | 139 |

### 2.4. Data analysis and visualization

To visualize the annual shift in soil GHG flux spatial patterns, contour maps of the cumulative soil GHG fluxes were created with ArcMap (Esri, Redlands, California, US), using inverse distance weighing (IDW) to spatially interpolate between datapoints. For this, the in-season cumulative flux was calculated for each measurement point with linear interpolation. To characterize the inter-annual stability of the spatial patterns and hotspots of the two gases across-year, the mean relative difference (MRD), and standard deviation of mean relative difference (SDRD) were calculated (Ashiq et al., 2021). The z-scores of MRD and SDRD values were calculated, and points with MRD z-score ≥ |1.0| and SDRD z-score ≤ 1.0 were considered consistent cold- or hotspots. Within each site and for each year, the z-scores of the cumulative GHG fluxes were also calculated to determine the hot/coldspots of that year (z-score ≥ 1). At Villa Grove site, point I10 had significantly high N_2_O flux values both years and skewed the scale of MRD values, so the square root of the N_2_O values were used for this site. All the figures except the contour map was created using *ggplot2* package (Wickham, 2016) in R environment (version 4.2.2) (R Core Team, 2019). A sensitivity analysis was conducted to determine how much spatial resolution affects the accuracy of the estimate of field-wide GHG flux. For this, the percent errors between the field-wide average cumulative CO_2_ and N_2_O fluxes of the subsampled datasets and the original dataset were calculated. At each subsample size, points were randomly selected without replacement, then the percent differences between the field-wide mean cumulative fluxes of the original dataset and the subsampled dataset were recorded. This was iterated a thousand times for each sample size, and the subsample size was divided by measurement area to find sampling points ha^−1^ as the unit to compare across sites and years. Bar-plots were created to visualize how variable the field-scale estimation can be along the spatial resolution.

## 3. Results and Discussion

### 3.1. Field-wide CO_2_ was comparable across years and sites

Overall, the field-scale spatial variability of the soil CO_2_ flux did not differ significantly across sites and between two years (Table 1), at the CV of 22% on average. At Bondville site, the field-scale mean cumulative CO_2_ flux was 1225 g m^−2^ (CV: 18%) in 2021 and 1224 g m^−2^ (CV: 23%) in 2022. At Pesotum site, the average cumulative CO_2_ flux was 1239 g m^−2^ (CV: 24%) in 2021 and 1148 g m^−2^ (CV: 21%) in 2022 (Table 1). At Villa Grove site, the average cumulative CO_2_ flux value was 1511 g m^−2^ (CV: 27%) in 2021 and 1690 g m^−2^ (CV: 22%) in 2022. These metrics suggest that the spatial variability of soil CO_2_ flux does not vary significantly among similar systems. Furthermore, the low spatial variability observed in our dataset further suggests that monitoring soil CO_2_ flux could be sufficiently done at lower spatial resolutions.

Likewise, the field-wide mean CO_2_ flux was also similar across sites and years, especially for the sites under conservation management under crop rotation between corn and soybean. These field-scale soil CO_2_ fluxes are comparable to past reports and are within the typical range for the Midwestern corn or soybean systems (Allaire et al., 2012; Ding et al., 2007; Hu et al., 2023). The mean cumulative flux was on average about 32% greater in the conventionally managed Villa Grove site. This agrees with past reports (Campbell et al., 2014) and likely owes to faster soil C turnover by higher C:N corn residues triggering positive priming effect (Mazzilli et al., 2014), and conventional tillage incorporating the fresh residues and soil organic matter while aerating the soil (Omonode et al., 2007; Ruis et al., 2022). Although this study cannot separate the effects of crop rotation and tillage, our results demonstrated that the combination of conventional tillage and continuous corn can increase the in-season soil CO_2_ compared to conservation practices.

### 3.2. The mean field-scale N_2_O flux and its spatial variability differed significantly by crop rotation and management

The spatial variability of soil N_2_O flux nearly doubled in the conventionally managed Villa Grove site compared to the two conservation management sites. The field-wide mean cumulative N_2_O flux at Bondville site was 155 mg m^−2^ (CV: 70%) in 2021, and 86 mg m^−2^ (CV: 94%) in 2022 (Table 1). Similarly, this was 172 mg m^−2^ (CV: 83%) in 2021, and 70 mg m^−2^ (CV: 63%) in 2022 for Pesotum site. Compared to these two sites, the field-wide mean cumulative N_2_O flux was substantially greater at Villa Grove site with 604 mg m^−2^ (CV: 135%) in 2021 and 785 mg m^−2^ (CV: 139%) in 2022. Meanwhile, the field-wide mean N_2_O flux was also greater in Villa Grove site, on average 4.2 times greater than those of Bondville and Pesotum sites during their corn year and 8.9 times than in their soybean year (Table 1). Likewise, the field-wide mean N_2_O flux doubled during the corn year for Bondville and Pesotum sites under conservation management.

Like CO_2_, these mean cumulative soil N_2_O fluxes were within those of the past reports in typical Midwestern corn and soybean systems (Bilen et al., 2022; Lawrence et al., 2021; Liu et al., 2017). Indeed, soil N_2_O flux is known to increase with corn cropping system in this region that typically receives high rates of N fertilizers (Ashiq et al., 2021; Jones et al., 2011; Morris et al., 2013; Tallec et al., 2019). Also, soil N_2_O increases under conventional tillage compared to conservation tillage, unless the soil is poorly drained (Bilen et al., 2022; Weidhuner et al., 2022). Thus, the continuous application of N fertilizers for corn monoculture in Villa Grove site would have synergized with the conventional tillage, leading to significantly greater field-scale N_2_O flux in this site compared to the other two sites, which also led to greater spatial variability (Bremner et al., 1981; Liu et al., 2017; Shcherbak et al., 2014). Our dataset demonstrated that compared to CO_2_, soil N_2_O flux is innately more spatially variable, and its spatial variability is highly sensitive to management like conventional tillage and N fertilizers, especially in corn cropping systems.

### 3.3. Contributions of CO_2_ and N_2_O hotspots to field-wide fluxes

The differences in spatial variability across sites and years also affected how much the GHG hotspots contributed to the field-wide emissions (Table 2). Averaged across years, each CO_2_ hotspot accounted for similar proportions of the field-wide CO_2_ flux, at 6%, 7%, and 6% in Bondville, Pesotum, and Villa Grove sites, respectively. These proportions were not significantly greater than those of each cold/intermediate spot at 4% (Bondville), 5% (Pesotum), and 4% (Villa Grove). On average, CO_2_ hotspots from 20%, 16%, and 13% of the total area contributed 26%, 22%, and 19% of the field-wide CO_2_ flux in Bondville, Pesotum, and Villa Grove sites, respectively. The small differences between hotspot contributions and those of non-hotspots reflected the small spatial variability (Table 1) and cumulative flux ranges of CO_2_ (Fig. 2).

**Figure 2.**
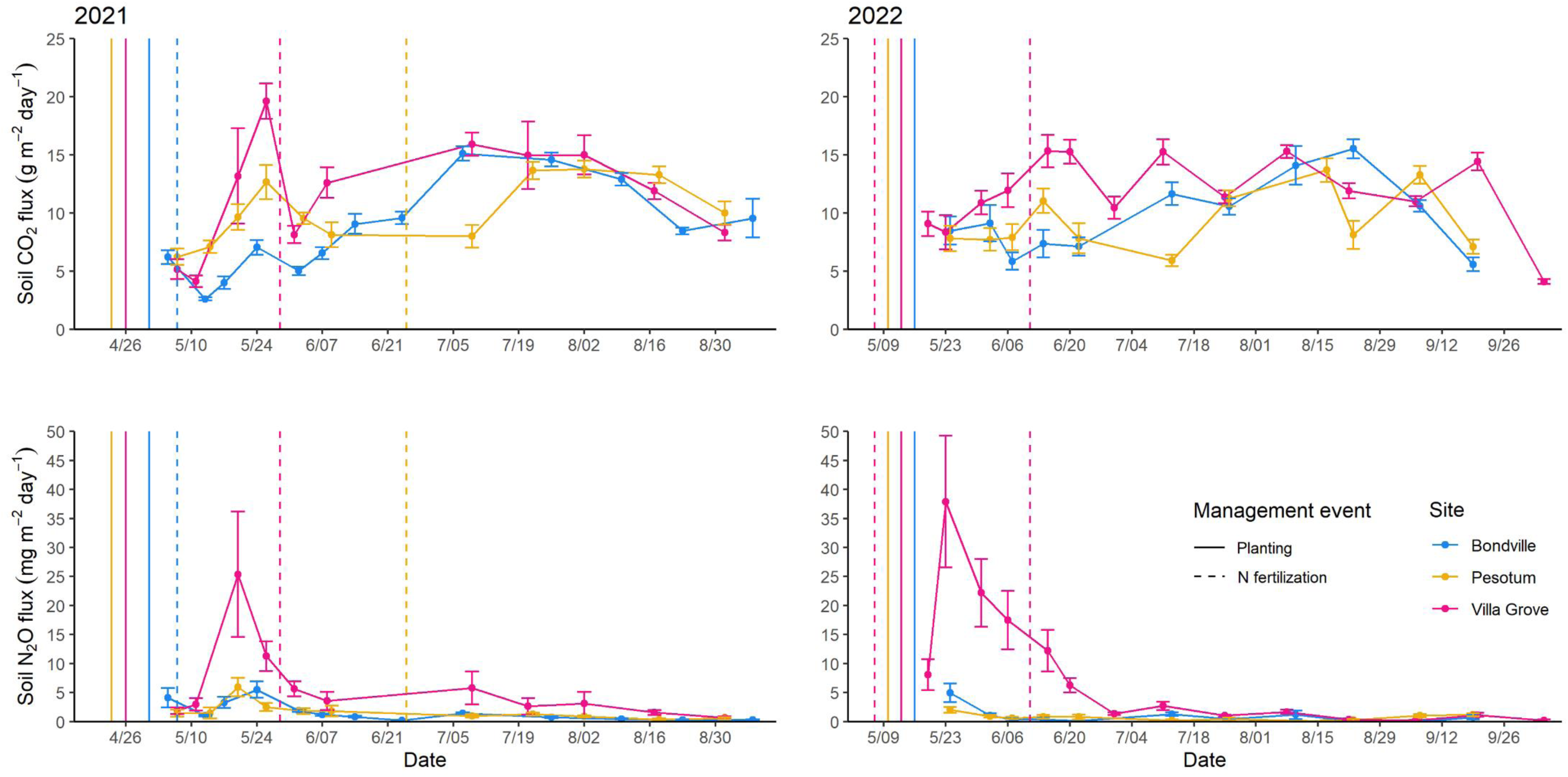
Timeseries of daily across-node averaged soil CO_2_ flux (top) and N_2_O flux (bottom) in 2021 (left) and 2022 (right) from each of the three sites (Bondville, Bondville; Pesotum, Pesotum; Villa Grove, Villa Grove). Error bars indicate standard error of the means. The vertical lines indicate major management events including planting and N fertilization.

**Table 2.** The mean contribution (%) to the field-wide cumulative soil GHG fluxes from each hotspot (z-score > 1), percent of the field the hotspots occupy, total contribution (%) to the field-wide flux, and the mean contribution (%) to the field-wide flux from each intermediate (z-score ≤ |1|) and coldspots (z-score < `−`1) for the in-season cumulative soil CO_2_ and N_2_O fluxes in Bondville, Pesotum, and Villa Grove sites in 2021 and 2022. Hotspots were considered as nodes where the z-score of the cumulative flux for that year was above 1.0.

| Site | Bondville |  | Pesotum |  | Villa Grove |  |
| --- | --- | --- | --- | --- | --- | --- |
| Year | 2021 | 2022 | 2021 | 2022 | 2021 | 2022 |
| Crop | Corn | Soybean | Corn | Soybean | Corn | Corn |
| Tillage | No-till | Reduced | Reduced | No-till | Conventional | Conventional |
| n | 40 | 16 | 19 | 19 | 18 | 40 |
| CO<sub>2</sub> |  |  |  |  |  |  |
| Mean hotspot contribution per node (%) | 3 | 8 | 8 | 7 | 9 | 4 |
| Hotspot area (%) | 15 | 25 | 16 | 16 | 11 | 15 |
| Total hotspot contribution (%) | 19 | 34 | 23 | 21 | 18 | 21 |
| Mean med/coldspot contribution per node (%) | 2 | 6 | 5 | 5 | 5 | 2 |
| N<sub>2</sub>O |  |  |  |  |  |  |
| Mean hotspot contribution per node (%) | 8 | 17 | 15 | 11 | 24 | 11 |
| Hotspot area (%) | 8 | 19 | 11 | 16 | 11 | 13 |
| Total hotspot contribution (%) | 23 | 50 | 30 | 32 | 49 | 54 |
| Mean med/coldspot contribution per node (%) | 2 | 4 | 4 | 4 | 3 | 1 |

Meanwhile, N_2_O hotspots contributed more disproportionately to the field-wide emissions than CO_2_. Each N_2_O hotspot accounted for 12%, 13%, and 18% of the field-wide fluxes in Bondville, Pesotum, and Villa Grove site, respectively, substantially greater than the average contributions from each cold/intermediate spot: 3% (Bondville), 4% (Pesotum), and 2% (Villa Grove) (Table 2). Indeed, N_2_O hotspots covering 13%, 13%, and 12% of the total area were responsible for 36%, 31%, and 51% of the field-wide fluxes in Bondville, Pesotum, and Villa Grove sites, respectively. Especially in Villa Grove site, hotspots that covered only 12% of the area contributed more than half of the field’s N_2_O emission. This level of disproportion with N_2_O hotspots has been observed in grazed grasslands, largely due to focused disposition of manure, which could be analogous to spatially heterogenous application of N fertilizers in the row crop systems (Cowan et al., 2015; Mason et al., 2017). A past study on a corn-soybean rotation system similar to Bondville and Pesotum sites reported that N_2_O hotspots (21% area) contributed 36% of the field-wide emission, relatively less disproportionate than our observations (Turner et al., 2016). However, this contrast could be the result of varying methods of defining hotspots, as they applied much more rigorous criterion (z-score ≥ 1.96) than ours, which emphasizes the need to standardize the definition of soil GHG hotspots to facilitate comparison between studies.

### 3.4. The inter-annual consistency of soil GHG flux spatial patterns and hotspots

The spatially interpolated contour maps of cumulative CO_2_ and N_2_O each year demonstrated that the spatial patterns, including the occurrences of hotspots, of these GHGs varied between the two years (Fig. 3). Our temporal sensitivity analysis at annual-scale (Fig. 4) demonstrated that while cumulative N_2_O fluxes are typically stable between years, hotspots will irregularly appear and only rarely will occur in consistent locations across years. Visualizing the relative GHG flux levels averaged across years (MRD), and their consistency between years (SDRD), the consistent hot/coldspots were indeed rare for both gases, with only one to three per site (Fig. 4).

**Figure 3.**
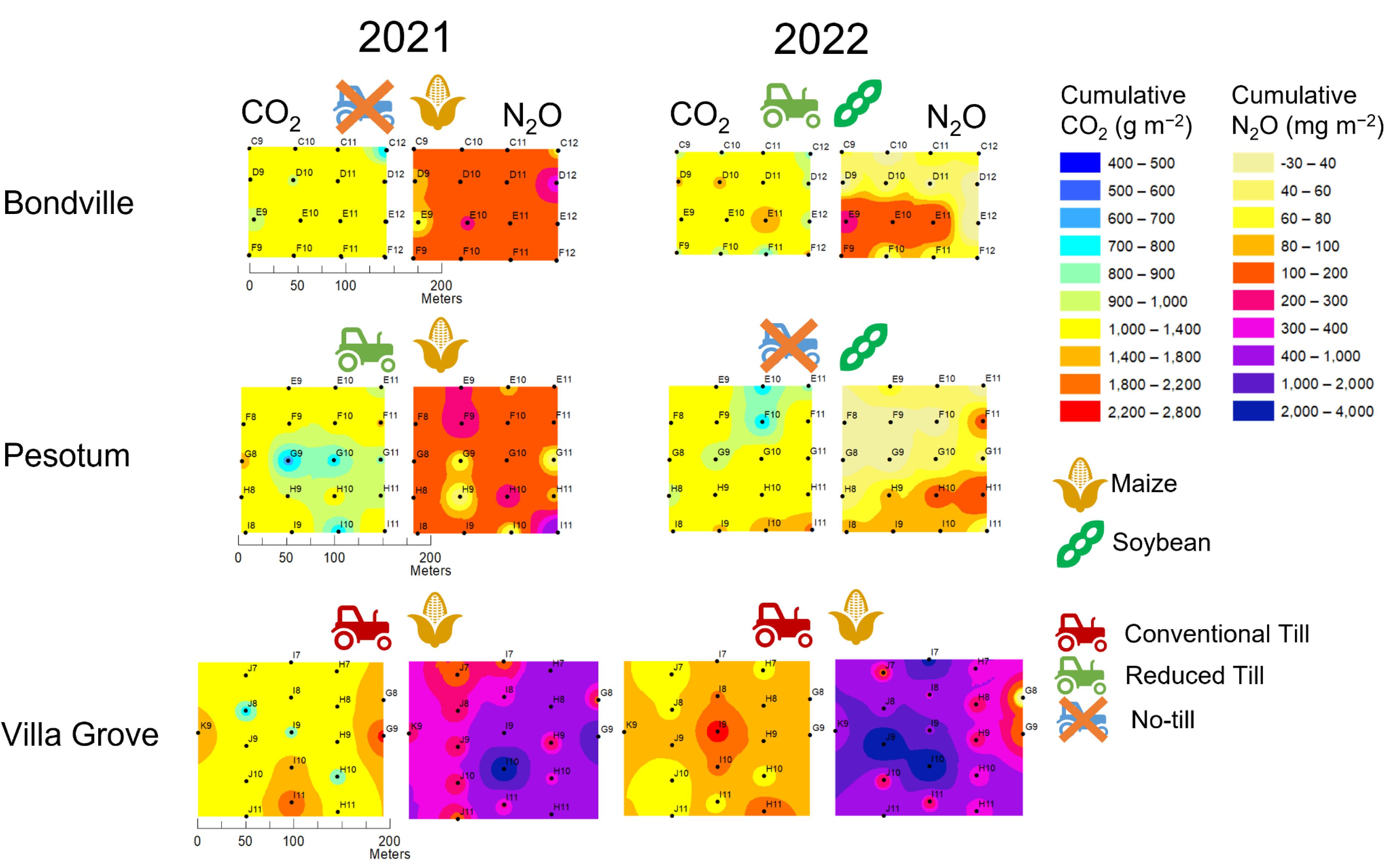
Inverse distance weighting (IDW) interpolation maps of in-season cumulative soil CO_2_ (left column of each year) and N_2_O (right column of each year) fluxes measured by manual chambers at the three sites (Bondville, Villa Grove, and Pesotum) for both years (2021 and 2022). The numbers of sampling points for Bondville, 2021, and Villa Grove, 2022, increased to forty but these extra nodes were not displayed to make comparison between years visually easier. Different icons for crops planted (maize and soybean) and tillage practices (conventional tillage, reduced tillage, and no-till) were placed above the maps to indicate the management of each site and year. Legends (top right) show the color codes of flux ranges designated to represent the distribution of the fluxes across sites and years.

**Figure 4.**
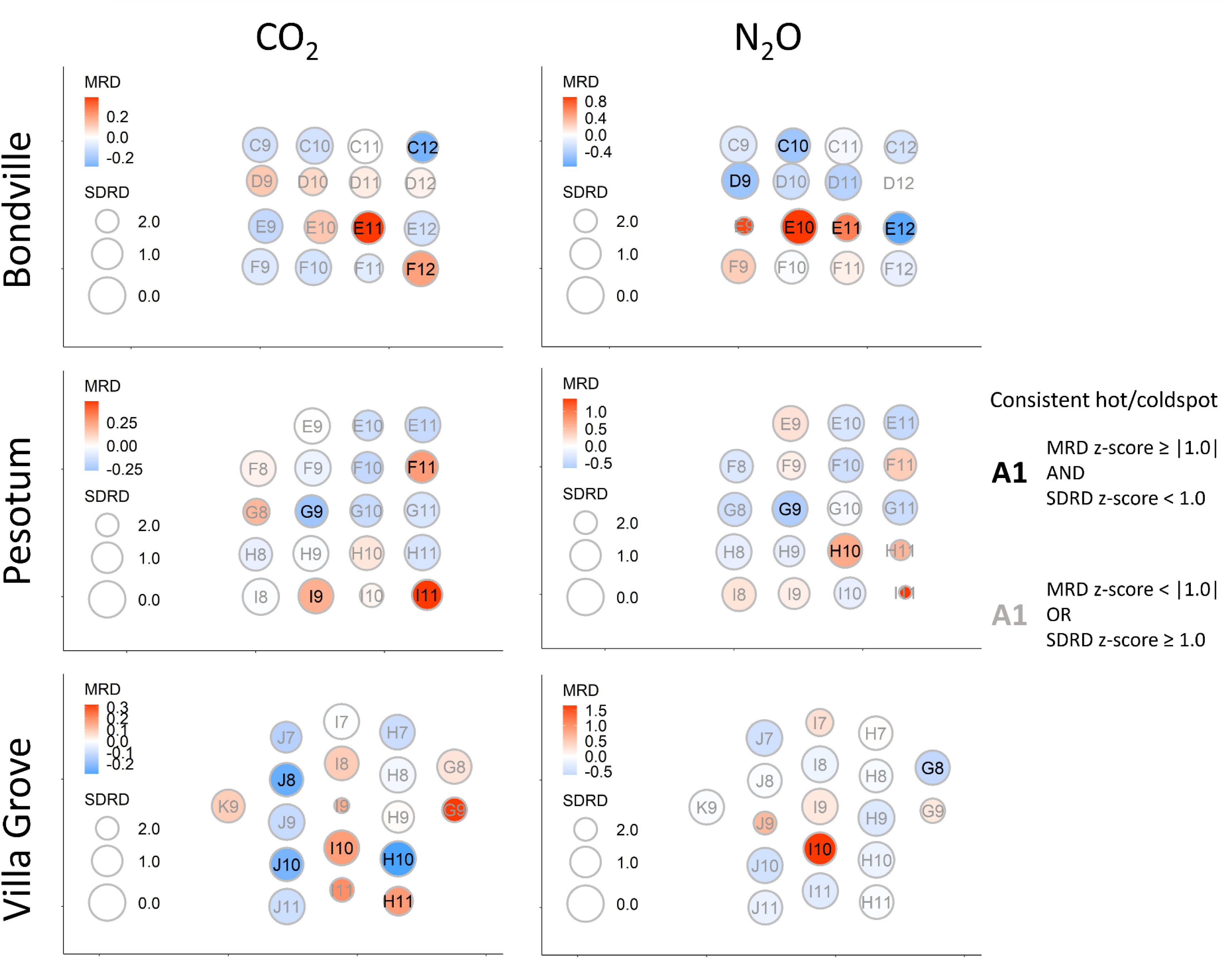
The inter-annual stability of cumulative soil CO_2_ (left column) and N_2_O (right column) fluxes for each of the three sites (Bondville, top; Pesotum, middle; Villa Grove, bottom). The letters followed by numbers indicate the sampling point ID and location along the longitude (X-axis) and latitude (Y-axis). The colors of the circles indicate the mean relative difference (MRD), the across-year average of the soil GHG flux values standardized within each site and year: red is relatively greater and blue relatively smaller. The size of the circle indicates the standard deviation of the mean relative difference (SDRD), with larger circle indicating low SDRD or greater stability between years and smaller circle indicating high SDRD or lower stability. Sampling point IDs in black indicate consistent hot or coldspots where Z-score of the MRD was greater than |1.0| and Z-score of the SDRD was smaller than 1.0. The sampling point IDs in grey indicate either points with intermediate fluxes or those that were inconsistent between years.

Meanwhile, measurement points with large difference in flux values between years were also uncommon. These highly variable points were typically hotspots in one of the years, which was most prominent in Villa Grove site like points J9, I7, and G9 where the cumulative flux values were more than ten-fold greater in one of the years (Figs. 3 and 4). A study of N_2_O flux from perennial grass systems over two years observed a greater number of consistent hotspots than this study, but this study was spatially smaller scale (each plot 0.4 ha) and how comparable perennial grass systems are with row crop systems is uncertain (Mason et al., 2017). A study conducted in Ontario, Canada, was similar to ours in both spatial and temporal scales (Ashiq et al., 2021). However, the authors composited the flux data throughout two growing-seasons and a winter fallow in-between to analyze the temporal stability. Hence, the authors demonstrated the spatiotemporal variation in N_2_O flux but not how it changes between years. Therefore, there are no comparable reports that currently exist to compare the inter-annual inconsistency in spatial patterns of soil GHG fluxes that we observed.

The inter-annual inconsistency that we observed in hotspots and spatial patterns of soil GHG fluxes elucidated a problem with the traditional methods to detect GHG hotspots at field-scale. One of the major motivations behind monitoring the spatial pattern of soil GHG flux over multiple years is to detect hotspots, to characterize them and ultimately use that information to mitigate them and reduce field-scale emissions. N_2_O flux is largely dictated by the interactions between soil nutrient availability (Nan et al., 2016; Shcherbak et al., 2014) and soil properties (Ball et al., 1997). At larger-scale, topographical gradients determine the soil properties that are relevant to N_2_O flux, like water-filled pore space and soil moisture, and they have been identified as useful predictors of N_2_O hotspots (Hall et al., 2018; Kravchenko et al., 2017; Turner et al., 2016). However, management practices, which are often spatially and inter-annually heterogenous, and their unpredictable coincidence with measurement equipment raise a question whether the spatial patterns truly shifted inter-annually, or the observed shift is simply a chance error. For example, only a couple of years of data cannot discern whether a sampling point was a true-positive hotspot in one year but not the next year due to its biogeochemical conditions, or simply because the N fertilizer was applied at the location of the measurement equipment in one year but not in the next. A multi-year dataset that can show a long-term pattern of each measurement point might overcome the errors caused by the spatiotemporally heterogeneous management effect and confidently identify a hotspot. However, collecting such long-term data is unrealistically resource and time consuming.

Our results emphasized that measurement campaigns for soil GHG fluxes should be designed according to the management practices and the goals of the campaigns. For example, broadcasted N fertilizers will be more spatially unpredictable than when knifed or side-dressed, as the latter is applied along the crop rows. For the latter case, equipment that can measure GHG flux across the within-row space may overcome the spatial heterogeneity that smaller equipment, as in this study, may overlook. If the goal is to identify hotspots and study their biogeochemical conditions, field experiments over areas that are small enough to rigorously control errors from spatially heterogeneous management practices and measurement locations may be more cost-effective. If the purpose of in-field monitoring is to estimate the field-wide emissions, measurements with high enough spatial resolution that can overcome the variability from both predictable and unpredictable sources with large sample sizes should suffice (Table 1).

### 3.5. Sensitivity analysis on soil GHG spatial variability to chamber measurement sampling density

Using the spatiotemporally high-resolution data collected from this study, we simulated lower-resolution datasets by randomly subsampling sampling points and compared them with the original dataset, thereby assessing the sensitivity of estimating field-wide emission to spatial resolution (Fig. 5). While a similar sensitivity analysis exists for temporal resolution (Lawrence et al., 2021), this has not been done for spatial resolution. Consistent with its low spatial variability, soil CO_2_ flux did not differ significantly between the simulated and original dataset even at low spatial resolution. Generally, the errors between all thousand iterations (excluding outliers) of simulated datasets and the original dataset for soil CO_2_ flux were within 25% of the mean field-wide flux of the original data when there were at least 1.6 sampling points per hectare, averaged across sites and years (Fig. 5). This result once again demonstrated that field-wide soil CO_2_ flux can be reliably estimated at low spatial resolution. In contrast, soil N_2_O flux required 5.6 sampling points ha^−1^, on average across sites and years, for the errors between the original and the simulated datasets to be within 25%. Low spatial resolution led to a bias where the estimated field-wide cumulative mean N_2_O flux was underestimated compared to the original dataset (Fig. 5). This resulted because randomly selected measurement points often did not include relatively rare N_2_O hotspots, thus underestimating the field-wide emission. Our simulations provide a reference to spatial resolution needed for a robust measurement campaign, with accurate estimates of field-scale N_2_O fluxes requiring much greater spatial resolution measurements than CO_2_.

**Figure 5.**
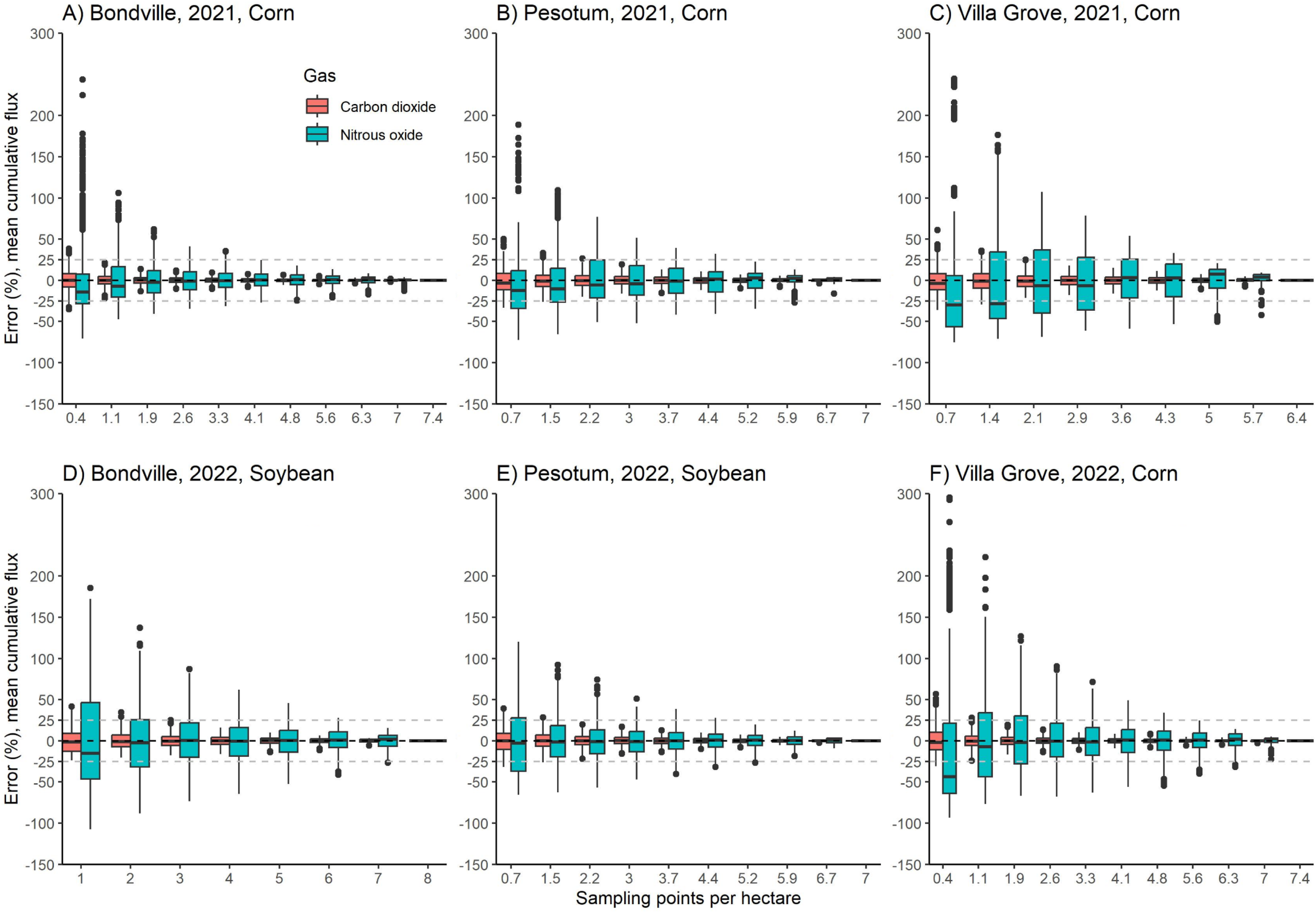
The percent errors between the mean field-wide cumulative soil CO_2_ (pink) and N_2_O fluxes (teal) of simulated datasets with lower spatial resolution and the original dataset for each year (rows) and sites (columns). The Y-axis shows the percent error between the simulated datasets and the actual dataset and X-axis denotes the spatial resolution of the simulated dataset as the number of randomly sampled points per hectare (actual area of data collection). A thousand iterations of simulated datasets were generated per spatial resolution. The box indicates the second and third quartiles and the top and low whiskers indicate the first and fourth quartiles. The thick horizontal line inside the box indicates median, and the dots indicate iterations whose percent errors were outliers.

## 4. Conclusion

This study showcased a rare multi-site, multiple years, full in-season, spatially high-resolution and field-scale dataset on the soil GHG fluxes of a major row crop system. We show that conventional practices like deep-chisel tillage and continuous corn receiving high-rate N fertilization might increase the field-scale emission and spatial variability of soil N_2_O flux. This highlights that not only will conventional practices increase N_2_O emission, but also compromise the ability to accurately monitor it. Conversely, soil CO_2_ flux was stable across sites and years, suggesting that monitoring soil CO_2_ flux will not require high spatial resolution measurement. Most soil GHG flux hotspots shifted between years, which we attribute to the unpredictability of observing true-positive hotspots due to spatially heterogenous management effects. Especially, fertilization locations among and within crop rows and their coincidence with the locations of measurement equipment may have affected where hotspots were observed, raising the question whether traditional field-scale monitoring method can confidently detect hotspots. A campaign practicing rigorously controlled field management and measurement procedures could yield a dataset that elucidates reliable spatial patterns of soil GHG flux but doing this at large spatial scale and resolution is impractical. Thus, field experiments over smaller areas could be the better approach to study the conditions and mechanisms behind N_2_O hotspots. Finally, our data suggests that if the purpose is to estimate field-scale emission, a grid deployment of chambers above a sufficient sample size, 1.6 and 5.6 measurement points ha^−1^ for CO_2_ and N_2_O, respectively, may provide reliable results.

## CRediT authorship contribution statement

**Nakian Kim:** Writing – Original Draft, Methodology, Visualization, Formal analysis, Investigation. **Chunhwa Jang**: Writing – Review & Editing, Investigation. **Kaiyu Guan**: Conceptualization, Methodology, Resources, Project administration, Funding acquisition**. DoKyoung Lee**: Conceptualization, Methodology, Resources, Project administration, Funding acquisition, Supervision, Writing – Review & Editing. **Evan DeLucia**: Conceptualization, Methodology, Resources, Project administration, Funding acquisition, Writing – Review & Editing. **Wendy H. Yang**: Conceptualization, Methodology, Resources, Project administration, Funding acquisition, Writing – Review & Editing.

## Declaration of competing interest

None

## Acknowledgements

The authors acknowledge that this work was supported by support from DOE ARPA-E SMARTFARM program DE-FOA-0001953. We thank Mary Marsh, Dana Landry, Marissa Chavez, Sunbong Jung, Jungwoo Lee, Kayla Vittore, and Nictor Namoi for their help in data collection and equipment maintenance. We also extend our thanks to William Eddy, Emily Stuchiner, and Ziliang Zhang for their help with methods development and data interpretation.

## Supplementary table legends

**Supplementary Table 1.** Detailed description and the dates of the management in the monitored sites.

## Supplementary figure legends

**Supplementary Figure 1.**
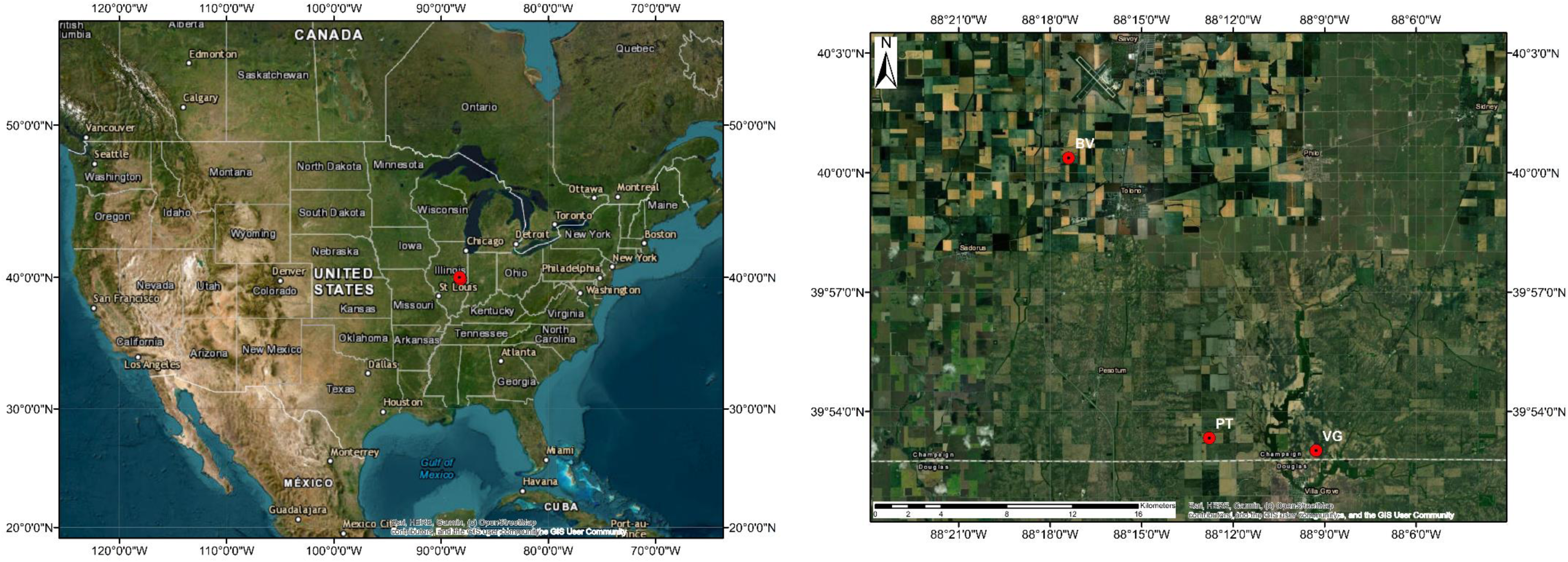
Satellite image of the locations (red pins) of the three experimental sites within Champaign County, IL, and the county’s location with the US.

## Notes

### Competing Interest Statement

This work has been supported by DOE ARPA-E SMARTFARM program DE-FOA-0001953

