## Supplementary figures and images for "Spatial variability of agricultural soil carbon dioxide and nitrous oxide fluxes: characterization and recommendations from spatially high-resolution, multi-year dataset"

### Supplementary figure 1

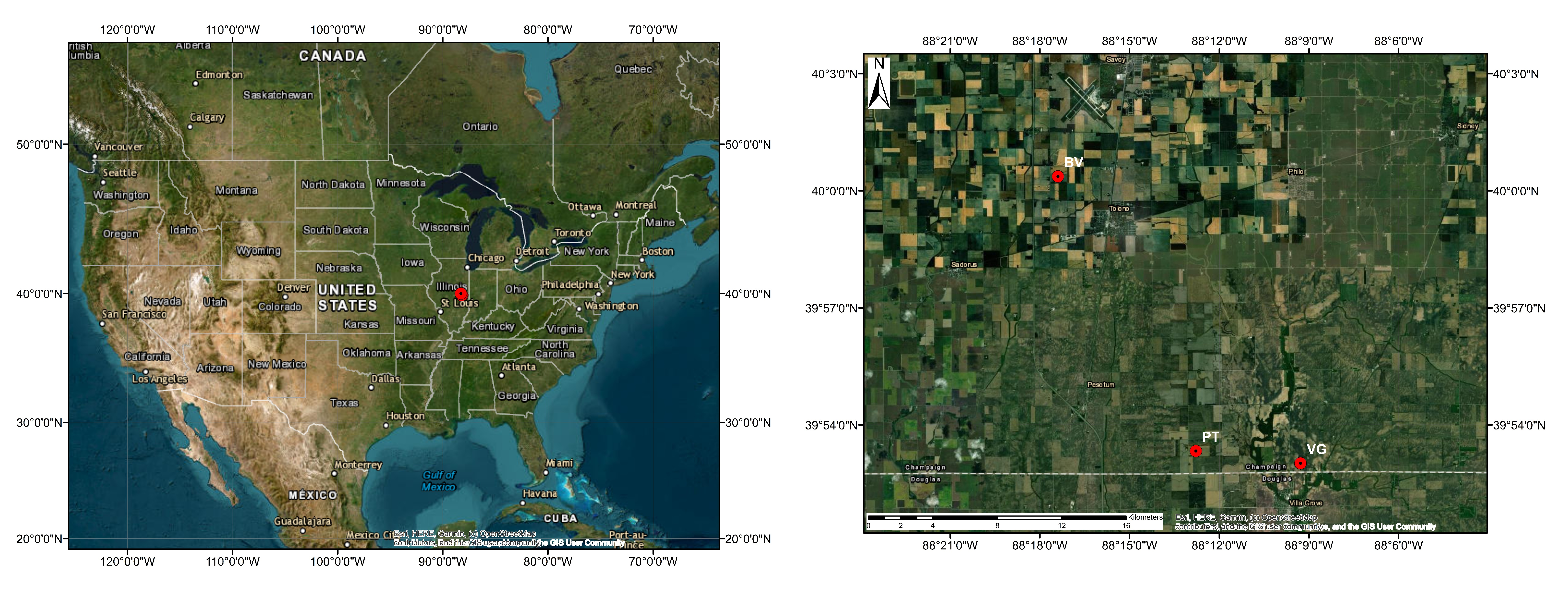
